# Faster attentional focusing in working memory instills better long-term memory

**DOI:** 10.1101/2024.03.25.586271

**Authors:** Sisi Wang, Freek van Ede

## Abstract

Working memory serves as a key gateway to the formation of lasting memories. While it is established how attentional focusing during working memory prioritizes internal representations for imminent tasks, the dynamic processes by which such internal focusing affects subsequent long-term memory remain underexplored. We developed a two-stage visual working-memory/long-term-memory task in which we cued visual objects during working memory and tracked the dynamics of attentional deployment through a recently uncovered gaze marker of internal focusing. Across two experiments, we found that attentional focusing during working memory benefits subsequent long-term memory for internally attended objects without a cost to unattended objects. Gaze biases associated with internal focusing revealed how this benefit was supported by the speed of attentional deployment, with faster attentional deployment predicting better subsequent memory. These results highlight how attentional focusing in working memory benefits long-term memory, and uncover the dynamic processes that instill such lasting benefits.

## Introduction

Working memory (WM) enables us to maintain relevant sensory information in the service of guiding behavior and has also been proposed as a crucial gateway to long-term memory (LTM) (Atkinson and Shiffrin, 1968; Baddeley, 2000; Cowan, 2008; Johnson, 1992; Jonides et al., 2008; Oberauer K and Hein L, 2012). Given the central significance of both WM and LTM for human cognition, extensive research has focused on elucidating how information held in WM is transferred to LTM, and identifying the factors that influence this transfer. For example, it has become well-established that longer maintenance and/or rehearsal of information in WM is associated with better subsequent LTM of this information (Aldridge and Crisp, 1982; Dark and Loftus, 1976; Darley and Glass, 1975; Hartshorne and Makovski, 2019; Souza and Oberauer, 2017). In a similar vein, it has been established that more attentive encoding of sensory information into WM (Bartsch et al., 2018; Craik and Lockhart, 1972; Hanslmayr et al., 2011; Khader et al., 2007, 2010; LaRocque et al., 2015; Ranganath et al., 2005; Sundby et al., 2019; Tozios and Fukuda, 2019; Voss and Paller, 2009) or retrieval of information from LTM back into WM (Carrier and Pashler, 1992; Chan and McDermott, 2007; Karpicke and Roediger, 2008; Roediger and Karpicke, 2006) can positively affect subsequent LTM.

Importantly, the fate of WM content is not only determined by the quality of encoding and the duration/quality of ensuing rehearsal, but also by selective-attention processes that are initiated during WM and that serve to prioritize specific mnemonic contents for guiding imminent behavior (Gazzaley and Nobre, 2012; Myers et al., 2017; Oberauer, 2019; Sahan et al., 2019; Souza and Oberauer, 2016; Van Ede and Nobre, 2023). To date, the majority of studies targeting such selective-attention processes during WM following so-called “retro-cues” (Griffin and Nobre, 2003) have focused on whether and how such cues improve performance on the ensuing WM task itself (for reviews, see Souza and Oberauer, 2016; Van Ede and Nobre, 2023). In contrast, whether and how attentional prioritization during WM in service of an upcoming WM task “spills over” into subsequent benefits on LTM remains less understood.

First, only a handful of retro-cue studies have studied the long-term consequences of selective object prioritization during WM (Fan and Turk-Browne, 2013; Hartshorne and Makovski, 2019; Jeanneret et al., 2023; LaRocque et al., 2015; Reaves et al., 2016; Sabo et al., 2024; Strunk et al., 2019; for complementary studies using “refreshing cues” see e.g., Bartsch et al., 2018; Johnson et al., 2002; Lintz and Johnson, 2021; Loaiza and Halse, 2019; McCabe, 2008). These studies have yielded only partially consistent results. While several studies have shown LTM benefits (Fan and Turk-Browne, 2013; Hartshorne and Makovski, 2019; Jeanneret et al., 2023; Reaves et al., 2016; Sabo et al., 2024; Strunk et al., 2019; Johnson et al., 2002; Lintz and Johnson, 2021; Loaiza and Halse, 2019; McCabe, 2008), others have not (Bartsch et al., 2018; LaRocque et al., 2015), or have suggested that retro-cue effects on LTM may be driven predominantly by a drop in performance for uncued (unattended) objects (Jeanneret et al., 2023), which may be explained simply by shorter retention in WM.

Second, a key open question pertains to the dynamic mechanisms that underlie the putative influence of attentional focusing during WM on subsequent LTM. Gaining insight into this pivotal question requires directly tracking the process of attentional focusing during WM. Recent work has uncovered that attentional focusing during visual WM can be robustly tracked through spatial biases in gaze (de Vries et al., 2023; Draschkow et al., 2022; Liu et al., 2022; van Ede et al., 2021; Van Ede et al., 2020; van Ede et al., 2019; Wang and van Ede, 2025, 2024). This opens unique opportunities for directly tracking how the process of spatially selective attentional focusing during visual WM affects subsequent visual LTM.

To examine whether and how attentional prioritization during visual WM affects LTM, we developed a two-stage visual WM-LTM task, while carefully controlling sensory exposure. Across two versions of this task, we investigated LTM consequences of selective focusing in visual WM, while also tracking such focusing directly through spatial biases in human gaze measurements. We demonstrate clear LTM benefits for visual objects that are cued during WM for guiding visual search, driven by a selective benefit for the cued (attended) object, without a cost to the other uncued (unattended) object. Critically, the continuous gaze marker revealed better subsequent LTM when attention was directed faster to the cued object during the preceding WM stage. These results highlight how attentional focusing during visual WM can have lasting benefits for visual LTM, and provide insight into the dynamic WM processes that instill such benefits.

## Results

Our results reveal whether and how attentional prioritization during visual WM affects LTM in three sequential steps. First, we establish the influence of retro-cueing during WM on subsequent LTM performance. Then, we establish a continuous time-resolved gaze marker as a marker of attentional allocation in WM which enables us to track the dynamic process of mnemonic prioritization following the retro-cue. Next, we link this dynamic time-resolved marker of attentional allocation in WM to subsequent LTM. For this we use the subsequent-memory-effect (i.e. SME), splitting trials by whether those objects that were cued during the WM stage were later remembered or later forgotten during the subsequent LTM stage. Finally, we address and rule out potential confounding factors that could underlie the observed link between retro-cueing during WM and subsequent LTM benefits.

### Attentional focusing during WM leads to better subsequent LTM recognition

In Experiment 1, each trial contained 1 cued (internally attended) and 1 uncued (internally unattended) memory object. One quarter of the search displays contained the memory target (the cued object), while the remaining three quarters of the search displays did not contain the target memory object. The uncued (unattended) object was never in the display. This yielded 4 conditions for investigating the effect of attentional focusing during WM on LTM recognition: cued-and-present, cued-and-absent, uncued-and-absent, and searched-only objects. The critical comparison here involves that between the cued and uncued memory objects in the target-absent-during-search conditions (i.e., cued-and-absent vs. uncued-and-absent). This enabled a direct comparison of the influence of attentional focusing during WM on subsequent LTM, while ensuring the same sensory exposure, given that neither object was repeated in the search display in these critical trials, even though one of them had been cued as the search target.

**Figure 1a** shows LTM recognition performance for all types of objects presented in the WM task. Our central a priori-defined comparison of interest revealed better LTM recognition accuracy for the cued-and-absent objects than for the recognition of the uncued-and-absent objects (*t*(24) = 4.714, *p* < 0.001, Cohen’s *d* = 0.986, Bonferroni corrected), with no significant difference of RT between the two (*t*(24) = -2.489, *p* = 0.091, Cohen’s *d* = -0.276). Given that the cued-and-absent and uncued-and-absent objects had the same sensory exposure (both only seen once during WM encoding) and the only difference between them is the selective attentional focusing on the cued objects during the WM delay, these results provide clear evidence that attentional focusing during WM for an upcoming WM task can have lasting consequences for subsequent LTM recognition.

**Figure 1.**
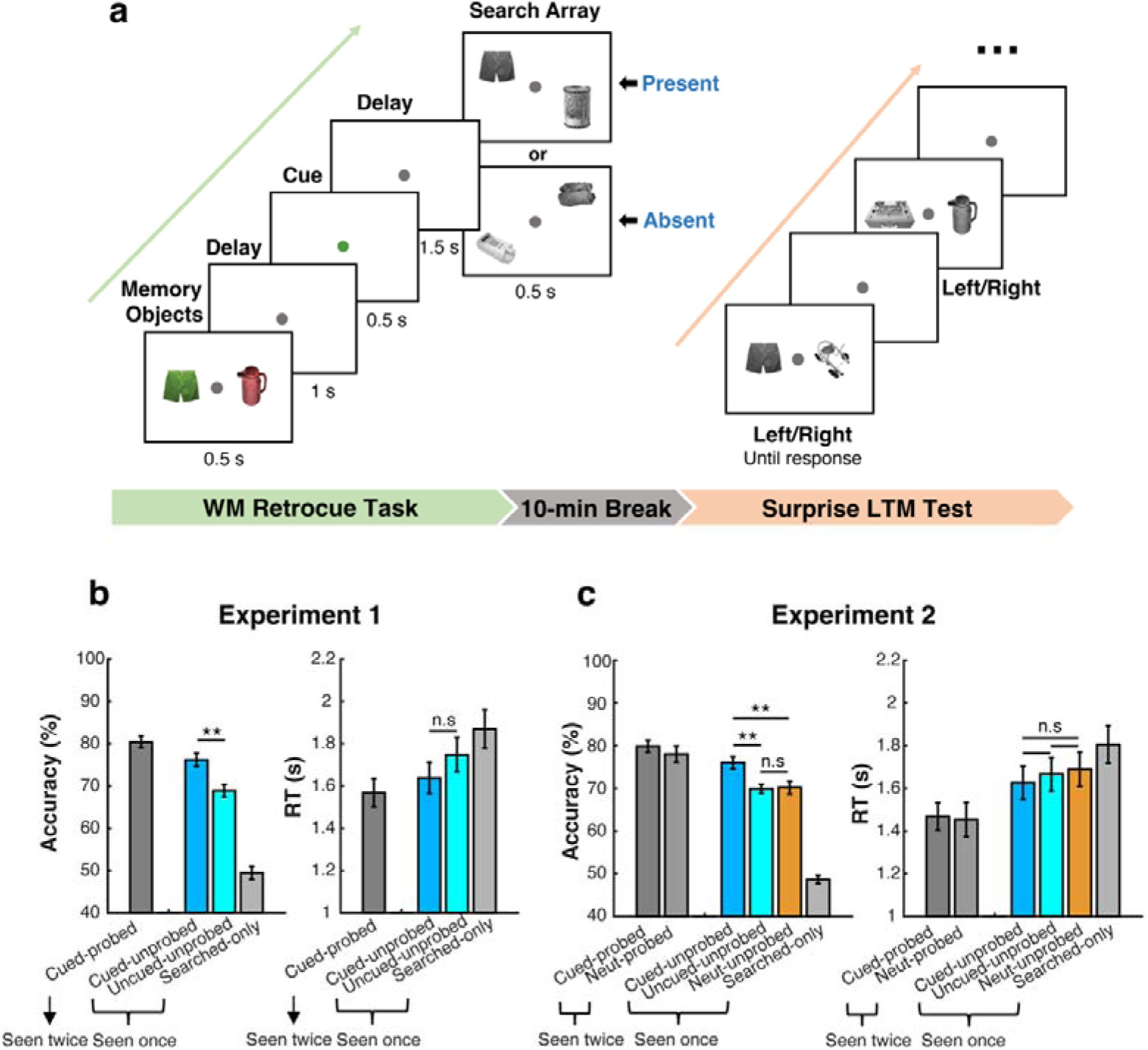
Experimental procedure and LTM recognition performance across experiments. **a**) The two-stage WM-LTM task procedure. In the visual WM task stage (left panel), participants memorized two visual objects, of which one could be cued (through its color) for an upcoming WM-guided search task. Upon the search display, participants were instructed to indicate the presence or absence of the cued object (or of both objects, in case of a neutral-cue trial that we included only in Experiment 2). Following a 10-minute break, they completed a surprise LTM test (right panel), where they needed to repeatedly indicate which one of the two objects they recognized from the preceding WM task. **b**) Performance on the surprise LTM recognition test as a function of attentional status of the objects during WM in Experiment 1 (containing exclusively informative cues). **c**) Performance on the surprise LTM recognition test as a function of attentional status of the objects during WM in Experiment 2 (containing both informative and neutral cues). Blue, cyan, and orange bars represent our key conditions from target-absent-during-search trials, consisting of objects that were cued-and-absent, uncued-and-absent, and neutrally cued-and-absent, respectively. Left and right panels show comparisons between recognition accuracy and response time (RT). Error bars represent ±1SEM. *, **, n.s represent significance levels *p* < 0.05, *p* < 0.01, and non-significant after Bonferroni correction. Note: only statistics of our main interest are highlighted in this figure. For a full overview of pairwise comparisons, see **Table S1**.

In addition to our central comparison of interest, we also found superior LTM recognition accuracy for the cued-and-present objects (likely owing to both repeated exposure and attentional focusing) compared to all other conditions (|*t*|s ≥ 2.747, *p*s ≤ 0.046, Cohen’s *d*s ≥ 0.575) and worst recognition for the searched-only objects that only served as fillers in the search display (|*t*|s ≥ 12.578, *p*s ≤ 0.001, Cohen’s *d*s ≥ 2.631). For a complete overview of all possible comparisons, see **Table S1**.

### The LTM benefit is driven by a facilitation of the attended memory object, not a cost of the unattended memory object

In Experiment 1, we observed a robust effect of attentional cueing during WM on subsequent LTM. However, because we only included 100% informative cues, we could not ascertain whether the effect on LTM was driven by a relative benefit for the cued (attended) object or a relative cost for the uncued (unattended) object. For example, it is possible that the uncued object showed reduced subsequent LTM performance because it could be dropped from WM midway through the trial (i.e., after the retro-cue), and was thus retained in WM for a shorter duration.

To address this possibility, we designed Experiment 2, in which we included neutral cues whose color did not match either object in WM. In these trials, participants were required to retain both objects until the search display and report whether either object was present (or, alternatively, whether both objects were absent).

The key difference between the informative and neutral cues is whether an attentional shift to one of the two memory objects is invited by the cue. Therefore, any significant difference in subsequent recognition accuracy between the focused/cued object and either object in neutral-cue trials must be attributed to whether attention is shifted toward the cued object during WM maintenance. Similarly, any significant difference in subsequent recognition accuracy between the unfocused/uncued object and objects in neutral-cue trials must be attributed to whether attention is shifted away from the uncued object after the cue.

By comparing the LTM performance for cued and uncued objects with the objects in trials with a neutral cue, we could resolve whether the cueing benefit on LTM was predominantly driven by a facilitation of the cued (attended) object or a cost to the uncued (unattended) object. Again, we focused on target-absent-during-search trials, where sensory exposure was equated.

First, results from Experiment 2 confirmed that participants used the informative cues during the WM task, as evidenced by superior search performance on the WM task in trials with informative vs. neutral cues (**Fig. S1**). This was the case for both accuracy and RT (Main effect of Cue Type, accuracy: *F*(1,49) = 29.668, *p* < 0.001, η*²_p_* = 0.377; RT: *F*(1,49) = 21.606, *p* < 0.001, η*²_p_*= 0.306).

Having confirmed the use of the cue during WM, we turned to our critical LTM results. As in Experiment 1, we observed reliable benefits on LTM recognition accuracy when comparing cued vs. uncued memory objects that were matched for sensory exposure (Fig. 1b, target-absent-during-search trials, *t*(49) = 4.268, *p* < 0.001, Cohen’s *d* = 0.621, Bonferroni corrected). Critically, these cueing effects were driven exclusively by a benefit of the cued (attended) object, as we found a clear LTM benefit for cued objects compared to neutral objects (Fig. 1b, t(49) = 4.052, *p* = 0.001, Cohen’s *d* = 0.589), while no statistically significant difference was found between uncued (unattended) and neutral objects (*t*(49) = - 0.216, *p* = 1.000, Cohen’s *d* = -0.031). A further Bayes Factor analysis showed that the current data were 6.259 times more evidence in favor of the null hypothesis than for the alternative hypothesis when comparing uncued (unattended) and neutral objects.

These data rule out that the cueing benefit on LTM is driven by a shorter retention duration of uncued (unattended) objects, and instead show how attentional focusing of the cued memory object brings about a benefit to subsequent LTM, compared to a control condition with the same retention duration but without attentional focusing during WM.

### Attentional deployment in WM can be tracked by directional biasing of gaze

Having established that attentional focusing during visual WM instills lasting benefits on visual LTM, we aimed to track the attentional deployment process in WM directly, to understand the mechanisms that underlie the influence of WM focusing on subsequent LTM.

Recently, it has been uncovered and reported how attentional shifts to visual objects in WM can be tracked by reliable spatial biases in eye movements (de Vries et al., 2023; Draschkow et al., 2022; Engbert and Kliegl, 2003; Hafed and Clark, 2002; Johansson and Johansson, 2014; Liu et al., 2022; Richardson and Spivey, 2000; Spivey and Geng, 2001; van Ede et al., 2021; Van Ede et al., 2020; van Ede et al., 2019; Wang and van Ede, 2025, 2024; Wynn et al., 2019). In our current data, we could also track attentional deployment through reliable spatial biases in saccade direction following the attentional retro-cue (Fig. 2), despite there being nothing to look at on the screen apart from the central fixation marker. Specifically, consistent with prior studies, we observed more saccades in the direction toward the memorized location of the cued object than in the opposite away direction – a finding that replicated across both experiments (Fig. 2a). A direct comparison of toward vs. away saccades (Fig. 2b) confirmed that these biases were highly reliable (permutation test, cluster-*p* = 0.002 for Experiment 1; cluster-*p* < 0.001 for Experiment 2). Moreover, by visualizing these toward vs. away biases as a function of saccade size (Fig. 2c), we observed that the spatial saccade biases included, to a considerable extent, saccades below 1 degree (within a range of microsaccades, Hafed et al., 2015; Martinez-Conde et al., 2013; Poletti, 2023; Rolfs, 2009), which was particularly clear in Experiment 2 for which we also had more data.

**Figure 2.**
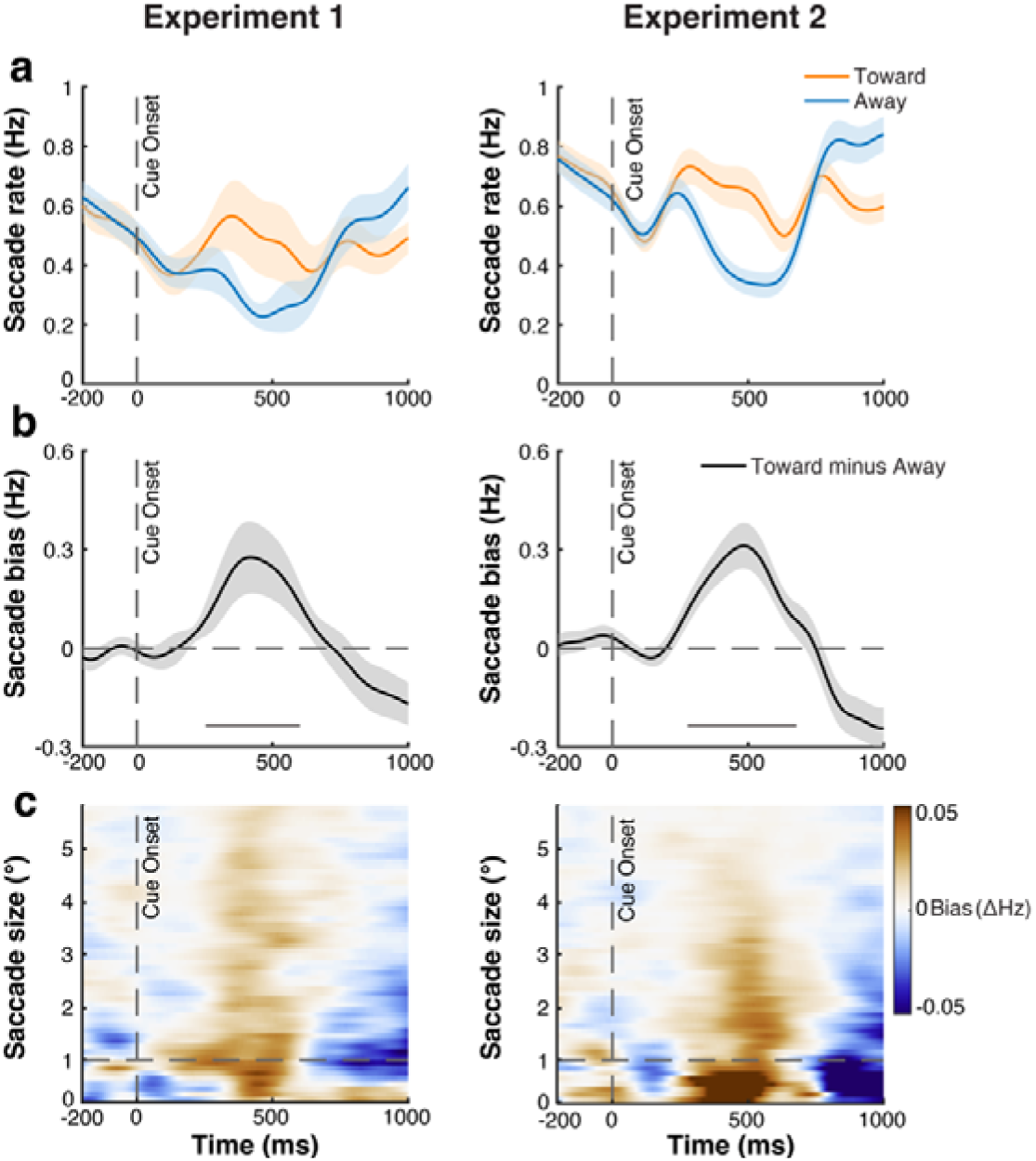
Spatial biasing of eye-movements tracks internal attentional focusing during WM in both experiments. **a**) Saccade rates toward (orange line) and away (blue line) from the memorized location of the cued object relative to cue onset in the WM task. **b**) The difference between toward and away saccades (i.e. the spatial bias). Horizontal grey lines above the x-axis represent significant clusters. **c**) Saccade rate effects (toward vs. away) as a function of saccade size, in degrees visual angle.

These findings replicate gaze biases during internal orienting of attention that have been reported previously, and show that such gaze biases also occur when directing attention to memorized images of real-world objects (see also Liu and van Ede, 2025) (as opposed to the simpler stimuli used in the majority of prior studies of this bias).

### Faster attentional deployment in WM is associated with better subsequent LTM

Having established the gaze marker of internal attention shifts in our WM task, we now turn to our central question regarding the dynamic processes by which internal attention shifts during WM confer LTM benefits. To this end, we adopted the logic of the ‘subsequent memory effect’ (SME; see e.g., Bartsch et al., 2018; Khader et al., 2007, 2010; LaRocque et al., 2015; Ranganath et al., 2005; Voss and Paller, 2009) by splitting the WM trials according to whether the cued objects were later correctly recognized (i.e., later-remembered) vs. incorrectly recognized (i.e., later-forgotten) in the LTM recognition test. While we found clear gaze biases following the attentional cue during WM, both for objects that were later-remembered and later-forgotten, we also observed faster gaze biases in trials in which cued objects were later-remembered (Fig. 3), consistent with an earlier—putatively more effective—attentional deployment.

**Figure 3.**
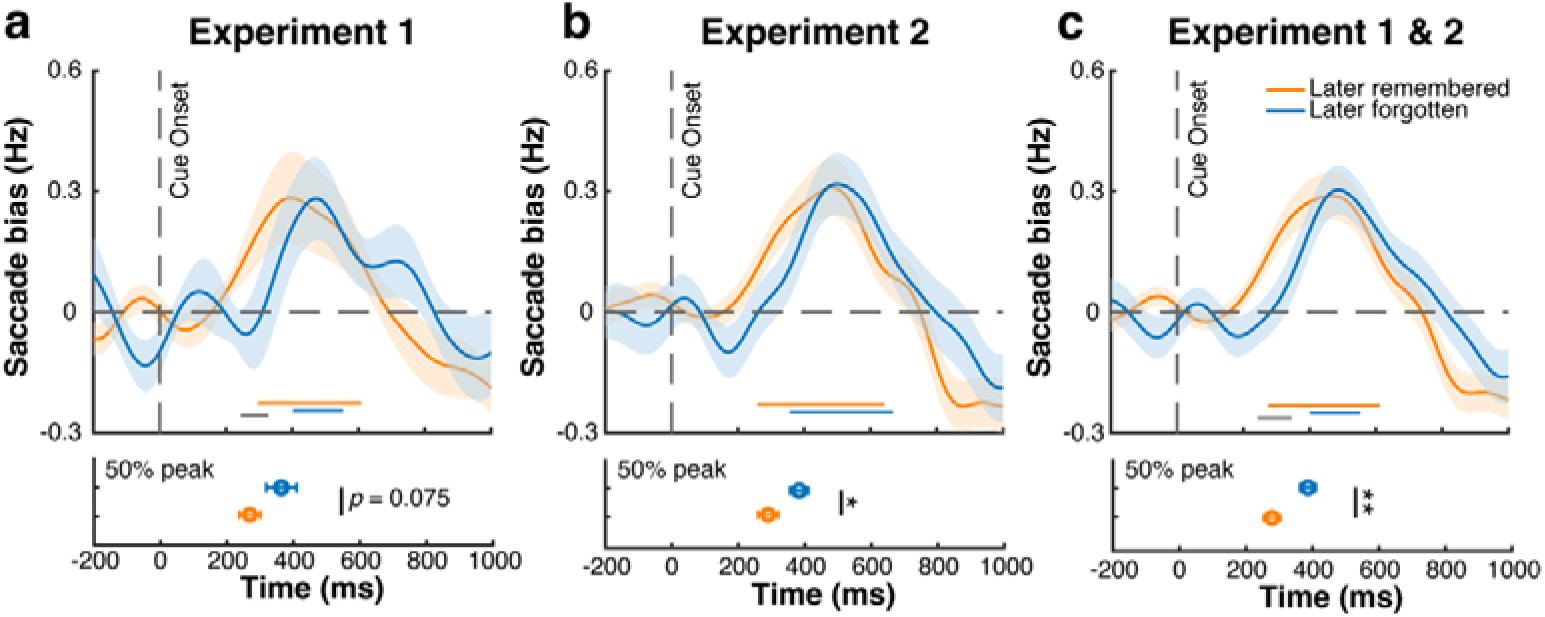
Faster attentional deployment in WM is associated with better subsequent LTM recognition. **a**) Top panel: spatial saccade bias (toward minus away) after the retro-cue during the WM task, for later-remembered and later-forgotten objects in Experiment 1. Bottom panel: latency of the saccade bias, defined as the 50% of the peak value The error bars represent standard errors obtained using a jackknife approach. **b**,**c**) Same as in panel **a**, but for Experiment 2 (panel b) and Experiment 1 and 2 combined (panel c). Orange and blue horizontal lines above the x-axis in top panels indicate significant clusters; grey horizontal lines represent significant difference clusters. Shading and error bars represent ±1SEM. *, ** represent significance level *p* < 0.05 and *p* < 0.01 after Bonferroni correction.

To quantify these differences, we directly compared attentional deployment—as tracked in our spatial gaze bias—for later-remembered and later-forgotten trials. We quantified gaze-bias latency as the first sample that reached 50% of the peak and quantified latency differences using a jackknife analysis (as in: Miller et al., 1998; Smulders, 2010).

As shown in Figure 3a**,b**, the pattern of delayed attentional deployment for later-forgotten memory objects was qualitatively similar in both experiments. When only considering the data from Experiment 1, the jackknife analysis on onset latency approximated significance (*t*(24) = -1.863, *p* = 0.075, Cohen’s *d* = -0.373) and reached significance after we equated trial numbers between later-remembered and later-forgotten trials, as we return to below. The same jackknife analysis in Experiment 2 – where we had more participants – confirmed a significantly faster gaze bias for later-remembered objects (*t*(49) = -2.212, *p* = 0.032, Cohen’s *d* = -0.313) that remained significant after matching trial numbers, as we return to below. Finally, for completeness, and to analyze this pattern with maximum sensitivity, we also combined the data from Experiment 1 and Experiment 2 (Fig. 3c). We could do so because the condition of interest here – the condition with an informative retro-cue – was identical in the two experiments. Interrogating these differences with increased sensitivity (Fig. 3c) corroborated a robust latency difference (*t*(74) = -3.349, *p* = 0.001, Cohen’s *d* = - 0.387). This latency difference was 108.2 ± 32.1 [M ± SE] ms.

In general, a larger number of trials contained objects that were later remembered vs. later forgotten. In Experiment 1, an average of 145.3 (SD = 13.2) and 46.7 (SD = 13.2) trials (cued objects) were subsequently remembered and forgotten in the LTM test; In Experiment 2, an average of 109.1 (SD = 13.1) and 34.9 (SD = 13.1) trials (cued objects) were subsequently remembered and forgotten in the LTM test. To rule out the possibility that our main gaze effect (of faster gaze bias during WM for later-remembered vs. later-forgotten objects) was driven by uneven trial numbers between these two conditions, we additionally ran a subsampling analysis where we randomly subsampled later-remembered objects with equal number of trials of the later-forgotten objects (repeating this subsampling 1000 times).

Our subsampling analysis, where we randomly subsampled later-remembered objects with equal number of trials of the later-forgotten objects, replicated our main results that later-remembered objects show faster gaze bias during WM than later-forgotten objects. Following this more careful analysis, the jackknife latency quantification reached significance in both Experiment 1 and Experiment 2 (**Fig. S2**, *t*(24) = -2.264, *p* = 0.033, Cohen’s *d* = - 0.453 and *t*(48) = -2.319, *p* = 0.025, Cohen’s *d* = -0.331 for Experiment 1 and Experiment 2, respectively). In addition to replicating the key effect of interest between Experiments 1 and 2, this shows that the reported differences between the saccade bias for later-remembered vs. later-forgotten objects cannot be attributed to uneven trial numbers between these trial-classes.

Having established an association between faster internal attentional deployment (as reflected in our gaze-bias marker) and better subsequent LTM performance, one might expect this effect to be particularly clear in individuals for whom attentional cueing in WM had a relatively large impact on LTM. To verify this, we split participants into two groups using a median split, based on their LTM benefit from attentional cueing during WM (i.e., the difference in LTM recognition accuracy between cued-and-absent objects and uncued-and-absent objects). Across the two groups, we observed an average of 125.9 (SD = 18.6) and 33.6 (SD = 10) trials (cued objects) that were subsequently remembered and forgotten for the large-LTM-benefit group, and 116.6 (SD = 24.5) and 43.8 (SD = 15.9) trials (cued objects) that were subsequently remembered and forgotten in the LTM test for the small-LTM-benefit group.

Consistent with our reasoning, and consistent across experiments (**Fig. S3**), faster attentional deployment for later-remembered vs. later-forgotten objects was particularly pronounced in those participants who showed the larger LTM benefit from WM cueing (Fig. 4). This pattern was verified by a two-way mixed repeated-measures ANOVA with a between-subjects factor of Group (small-LTM-benefit participants vs. large-LTM-benefit participants) and a within-subjects factor of the Subsequent Memory Effect (i.e. SME: later-remembered vs. later-forgotten) on jackknife-adjusted onset latencies (Smulders, 2010). The ANOVA revealed a significant main effect of SME (*F*(1,73) = 18.163, *p* < 0.001, η*²_p_* = 0.199) and an interaction between Group and SME (*F*(1,73) = 4.825, *p* = 0.031, η*²_p_*= 0.062). Post-hoc t-tests revealed that the interaction was driven by a clear SME effect in the large-LTM-benefit participants (*t*(36) = -4.537, *p* < 0.001, Cohen’s *d* = -0.920) that was significantly weaker (and not significant) in the small-LTM-benefit group (*t*(37) = -1.470, *p* = 0.875, Cohen’s *d* = -0.294), even if the trend was in the same direction in this group of participants.

**Figure 4.**
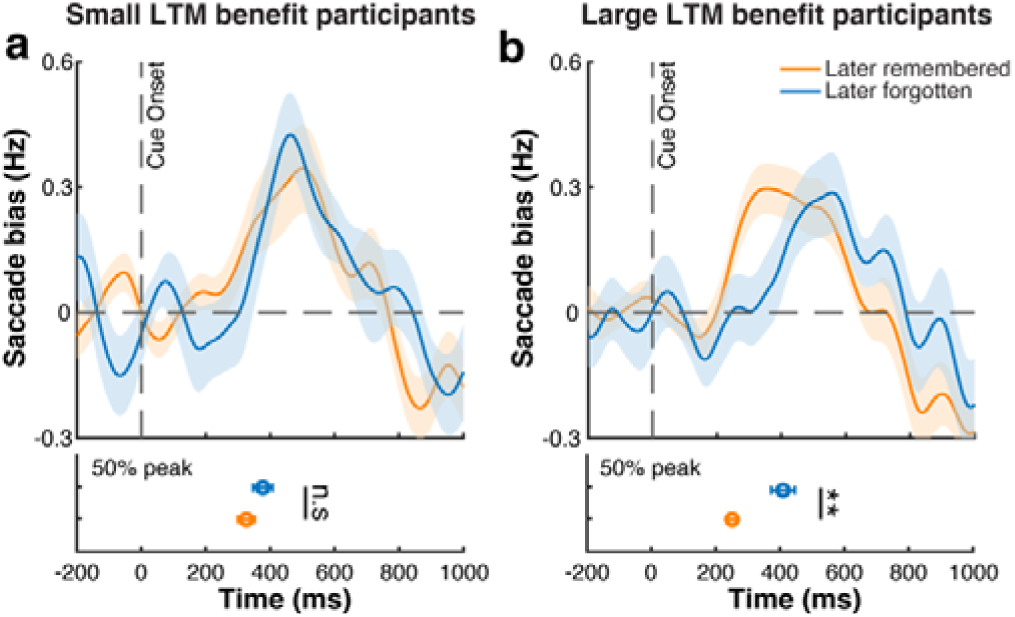
The latency of attentional deployment in WM predicts subsequent LTM particularly for individuals with larger LTM benefit. **a**) Gaze bias for later-remembered and later-forgotten trials in the subset of participants with relatively little benefit of cueing during WM on subsequent LTM. Conventions as in Figure 3. **b**) Gaze bias for later-remembered and later-forgotten trials in the subset of participants with relatively large benefit of cueing during WM on subsequent LTM. Shading and error bars represent ±1SEM. *, **, n.s represent significance levels p < 0.05, p < 0.01, and non-significant after Bonferroni correction. Note: to maximum the statistical power of this group analysis, the plots illustrate the merged dataset of Experiment 1 & 2. We found similar patterns across both experiments, as shown in **Figure S3**.

Together, these results provide evidence (from both the across-trial and the across-participant levels) that the speed (but not necessarily the strength) of attentional deployment within WM predicts the influence of mnemonic focusing on subsequent LTM performance. Faster attentional deployment during WM is associated with better subsequent LTM recognition, and this effect is particularly pronounced in those individuals who show the strongest influence of WM focusing on subsequent LTM.

Note how these findings help to argue against non-cue-specific explanations for the observed subsequent memory effect, such as generic fluctuations in alertness, arousal, or motivation at encoding (that may drive both better LTM and faster gaze bias after the retro-cue). Such generic influences would be expected to apply similarly to all participants, while our finding is particularly pronounced in those participants who benefitted a lot from the retro-cue. Accordingly, these data promote a “cue-specific” interpretation of the reported subsequent memory effect. Further support for a cue-specific interpretation comes from the observation that the latency difference is specific to the cued object (**Fig. S4**).

### The observed relation between the timing of attentional deployment and subsequent LTM is *not* driven by object memorability nor by time-on-task

We reported how the speed of attentional deployment in WM – as reflected in the reported gaze bias after the retro-cue – predicted subsequent LTM performance. This suggests that the speed (or, tentatively, the efficacy) of attentional deployment can have lasting consequences on memory. However, besides variability in the attentional process itself, may there be other, more trivial, generic explanations that can account for these key results? Below we consider, and rule out, two such viable alternative accounts.

#### Object memorability

First, it could be that some of our objects are simply more memorable, and that these more memorable objects are somehow also associated with a faster gaze bias when cued for selection during WM. To assess this first possibility, we estimated the memorability of all objects by sorting them by the overall correct identification rate during the LTM test (collapsed across our full sample of participants and combining both experiments for maximum sensitivity). To assess whether this measure of overall object memorability could account for latency differences in the gaze bias, we split the objects into high and low memorability (based on a median split) and assessed the time courses of the reported gaze bias as a function of memorability condition (Fig. 5a and **Fig. S5a**). This revealed no evidence for a statistically significant difference in the latency of the gaze bias when attention was directed to objects with generic low or high memorability (*t*(74) = -0.493, *p* = 0.624, Cohen’s *d* = -0.057). A further Bayes Factor analysis showed that the current data were 7.000 times more evidence in favor of the null hypothesis than for the alternative hypothesis.

**Figure 5.**
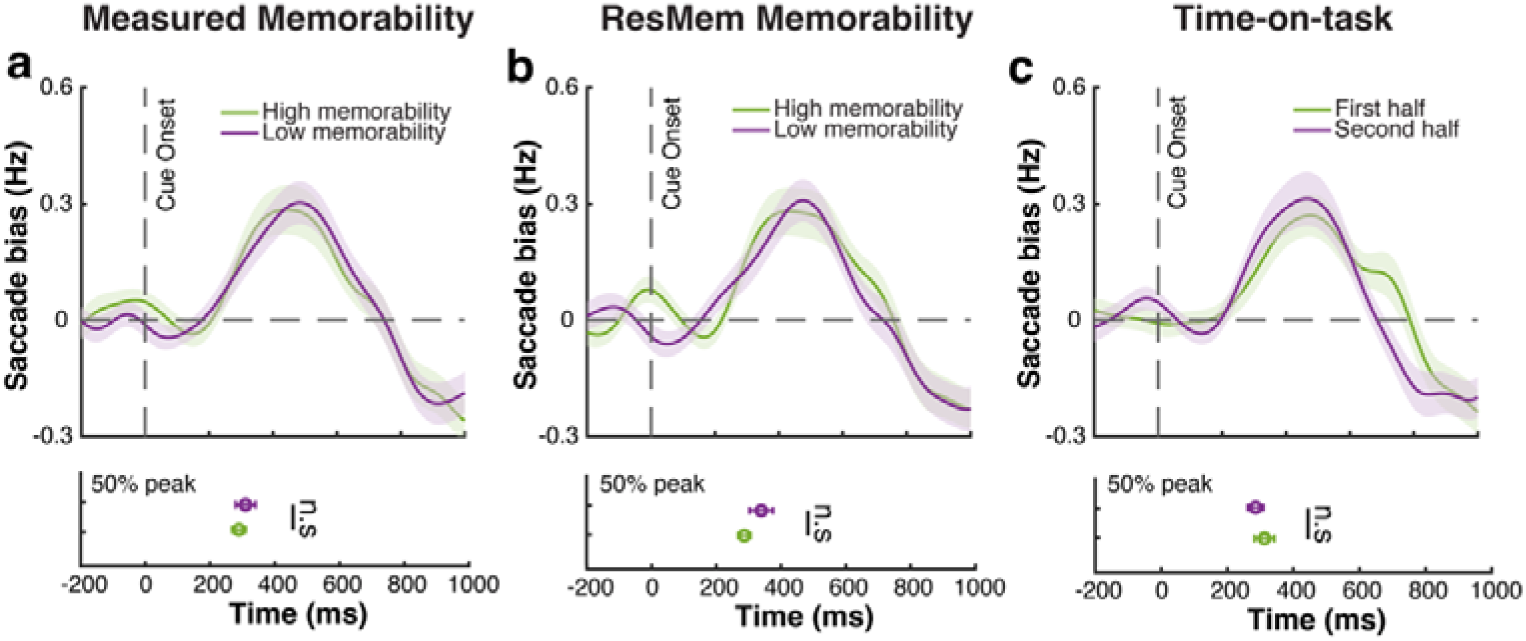
The observed relation between the timing of attentional deployment and subsequent LTM is not driven by object memorability nor by time-on-task. **a**) Spatial saccade bias after the informative cue during the WM task for cued objects with high (green) and low (purple) general memorability. Memorability was estimated from LTM performance from the current experiments, and objects were sorted using a median split. **b**) As panel a, but quantifying object memorability using ResMem (Needell and Bainbridge, 2022). **c**) Spatial saccade bias for objects presented in the first-half (green lines) vs. objects presented in the second-half (purple lines) of the WM session. Bottom panels and all other conventions as in preceding figures. n.s represents non-significant after Bonferroni correction. Note: to increase the statistical power, the plots illustrate the merged dataset from Experiments 1 & 2. Similar patterns were observed across both experiments, as shown in **Supplemental Fig. S4**.

In addition to our above approach, where we defined object memorability empirically based on our own LTM performance data, we also quantified object memorability using a state-of-the-art machine-learning model for predicting the intrinsic memorability of an image—ResMem (Needell and Bainbridge, 2022). We again found no evidence for a statistically significant difference in the latency of the gaze bias between objects with either low or high memorability, after having quantified memorability this way (Fig. 5b and **Fig. S5b**), (*t*(74) = - 1.311, *p* = 0.194, Cohen’s *d* = -0.151). A further Bayes Factor analysis showed that the current data were 3.465 times more evidence in favor of the null hypothesis than for the alternative hypothesis.

Together, these results suggest that the observed influence of the latency of attentional deployment during WM on subsequent LTM is unlikely driven by a difference in the memorability of the visual objects themselves.

#### Time-on-task

Second, we considered time-on-task during the WM task as another potential factor of the observed link between gaze bias latency and subsequent LTM performance. For example, gaze bias may become progressively faster during the WM task, and later trials in the WM may also have better LTM performance because they are more recent. To assess this second possibility, we also split our trials into the first-half vs. the second-half of trials in the WM task (Fig. 5c and **Fig. S5c**). Again, we found no evidence that time-on-task could account for the observed difference in latency of the gaze bias during the WM task (*t*_(74)_ = 0.821, *p* = 0.414, Cohen’s *d* = 0.095). A further Bayes Factor analysis showed that the current data were 5.690 times more evidence in favor of the null hypothesis than for the alternative hypothesis.

For complementary analyses and results of more fine-grained control analysis of Memorability and Time-on-task, see supplemental **Figure S6** and **S7**. Instead of relying on a median split, we additionally split all objects into 3 and 4 groups with different ranks of memorability from high to low, and split all trials into three and four sessions from early to late. These tertile- and quartile-split analyses, like our median-split analyses, showed a similar lack of any obvious modulation of our gaze bias by object memorability and time-on-task.

These additional analyses suggest that the reported effect of the speed of attentional deployment in WM — as tracked in the gaze bias — are not simply influenced by the objects themselves, nor by time on the WM task (and our data further speak against generic variability at encoding, as we considered in the preceding section). Instead, this central result likely reflects variability in the process of directing attention to internal WM contents, with consequences for LTM.

## Discussion

Our data reveal how focusing attention on visual objects held in WM for an upcoming WM task instills a lasting benefit on visual LTM recognition. We show that this is driven by a benefit to internally attended objects without a cost to unattended objects. In addition, we here uniquely tracked the process that supports this LTM benefit, capitalizing on spatial biases in eye movements that track focusing during visual working memory with high temporal resolution. This revealed that faster (not necessarily stronger) attentional deployment during WM is associated with better subsequent LTM for the attended WM object. These findings advance our understanding of the dynamic processes that underlie the influence of attentional focusing in WM on subsequent LTM.

### Attentional focusing during WM benefits subsequent LTM performance

Our behavioral performance data complements and reinforces several prior studies investigating the impact of attentional focusing during WM maintenance on subsequent LTM following either *retro-cues* (Fan and Turk-Browne, 2013; Hartshorne and Makovski, 2019; Jeanneret et al., 2023; LaRocque et al., 2015; Reaves et al., 2016; Sabo et al., 2024; Strunk et al., 2019) or *refreshing cues* (Bartsch et al., 2018; Johnson et al., 2002; Lintz and Johnson, 2021; Loaiza and Halse, 2019; McCabe, 2008). Like other prior work using retro-cues, our attentional manipulation involved focusing on specific memory objects in service of an upcoming WM task. Despite the fact that participants did not yet know they would later be tested for their LTM recognition of the objects, we found clear consequences of this type of WM focusing on subsequent LTM. By adding neutral cues, we could further show that this focusing operation engaged a benefit for the focused memory object, without incurring a cost to the other object in WM that could be dropped after the cue (cf. Astle et al., 2012; Gözenman et al., 2014; Gressmann and Janczyk, 2016; Griffin and Nobre, 2003; Gunseli et al., 2015; Oberauer, 2001; Pertzov et al., 2013; van Moorselaar et al., 2015; Williams et al., 2013). This is important because a cost to the uncued/unattended object (as reported in Jeanneret et al., 2023) could simply be explained by a shorter maintenance duration in WM (Aldridge and Crisp, 1982; Hartshorne and Makovski, 2019; Souza and Oberauer, 2017).

Our observed LTM benefits cannot be attributed to differential sensory processing, nor to differential sensory exposure. First, during encoding, participants could not yet know which of the two objects would become cued. Hence, sensory encoding was always equated between our attentional-cueing conditions. Further, note how this also separates our findings from ample prior work focusing on attentional influences on LTM encoding (Bartsch et al., 2018; Craik and Lockhart, 1972; Hanslmayr et al., 2011; Khader et al., 2007, 2010; LaRocque et al., 2015; Ranganath et al., 2005; Sundby et al., 2019; Tozios and Fukuda, 2019; Voss and Paller, 2009). Second, we made sure to include WM trials in which the cued memory objects were never repeated in the search display (requiring an ‘absent’ report). In these critical target-absent-from-search trials—that were randomly interleaved with target-present trials—participants used the retro-cue to prepare the search for the cued object, but the attended memory object never reappeared during the search. Thus, not only was sensory processing at encoding equated between cued/uncued/neutral trials, but so was total sensory exposure (all objects seen once) when considering these critical target-absent-during-search trials.

### Gaze patterns reveal how attentional focusing during WM influences subsequent LTM

A central novelty of our study is our focus on tracking the mechanisms by which attentional focusing during WM influences subsequent LTM. To provide new insight into this key underexplored question, we here took advantage of a recently established index of attentional deployment within the spatial layout of visual WM – spatial biases in gaze (de Vries et al., 2023; Draschkow et al., 2022; Liu et al., 2022; van Ede et al., 2021; Van Ede et al., 2020; van Ede et al., 2019; Wang and van Ede, 2025, 2024; for complementary findings, see also Engbert and Kliegl, 2003; Hafed and Clark, 2002; Johansson and Johansson, 2014; Richardson and Spivey, 2000; Spivey and Geng, 2001; Wynn et al., 2019). We first established that this gaze marker also tracked attentional focusing in WM in our case with images of real-world objects (see also Liu and van Ede, 2025). Having established this, we adopted the logic of the subsequent memory effect (SME, Bartsch et al., 2018; Hanslmayr et al., 2011; Khader et al., 2007, 2010; LaRocque et al., 2015; Ranganath et al., 2005; Voss and Paller, 2009) to address whether and how this marker of attentional deployment following the retro-cue predicted whether participants subsequently remembered vs. forgotten the cued objects. This revealed that the speed of attentional deployment during WM—not the degree of attentional deployment per se—predicts subsequent LTM.

An important question pertains to why faster attentional focusing may instill better long-term memory. We see at least two explanations. First, when shifting internal attention earlier, the attended object will be longer in the focus of attention until the search array appears. Being longer ‘in focus’ may confer a larger benefit. Though we are unable to directly track the duration of deployed attention with our gaze index (which likely reflects initial spatial orienting in working memory without necessarily also tracking the continued prioritisation of the cued object), future research using complementary measures, such as neural recordings, that enable to directly track the maintained memory content and its moment-to-moment priority status could provide more direct evidence for this mechanism. Second, faster internal attentional deployment may signal a more alert mindset during the retro-cue which may foster more effective attentional selection and prioritization of the cued object, in turn boosting memory consolidation. Delineating these alternatives and the precise neurobiological mechanisms underlying our findings represents a fruitful line of research opened by our results.

### Attentional focusing during WM vs. the testing effect

At face value, our data may appear reminiscent of other established findings from the LTM literature, such as the ‘testing effect’, whereby elements that are retrieved from LTM as part of a test show better subsequent LTM (in comparison to elements that are merely studied for longer, e.g., Carrier and Pashler, 1992; Chan and McDermott, 2007; Karpicke and Roediger, 2008; Roediger and Karpicke, 2006). While one can conceive our findings after the retro-cue as a form of ‘retrieval’, there are key differences with the aforementioned literature. First, our retro-cue prompted the selection of an object from WM, not from LTM. Moreover, it prompted a selection between one out of two possible objects, rather than one out of an almost infinite number of objects in LTM. For this reason, we have described our results from the perspective of attentional focusing in WM, consistent with a large body of research from the WM literature (Gazzaley and Nobre, 2012; Griffin and Nobre, 2003; Myers et al., 2017; Oberauer, 2019; Sahan et al., 2019; Souza and Oberauer, 2016; Van Ede and Nobre, 2023). By studying such selective focusing in the context of a WM task with visual objects, we were also in a position to directly track such focusing through spatial-attentional biases in gaze, yielding key novel insight into the ways by which such attentional focusing during WM affects LTM.

### Limitations and future directions

A major limitation of the subsequent-memory-effect approach we adopted here is its correlational nature. While we observed that the speed of attentional deployment following the cue predicted subsequent LTM, we cannot be sure that the speed of attentional deployment *causes* better subsequent LTM. Even so, we address and rule out viable alternative explanations for this observed effect, ruling out object memorability and time-on-task. Having ruled out these variables, we tentatively posit that our findings reflect genuine temporal variability in the process of directing attention to internal WM contents, with LTM consequences. Our findings from a secondary analysis in which we split participants by the size of their LTM benefit (Fig. 4) further corroborated this cue-specific interpretation.

We here defined WM and LTM by their respective stage in our task. At the same time, we note how even at the WM stage, the images of real-world objects that we used likely received contributions from LTM associations (Bartsch and Shepherdson, 2021; Brady and Alvarez, 2011; Curby et al., 2009; Park and Awh, 2025). This will generally be the case during everyday visual WM when holding in mind real-world objects, such as when we turn around and remember what is behind us. We show how attentional deployment at this WM stage has consequences for what we later remember in a subsequent LTM test. In future work, it will be interesting to also manipulate the degree to which the objects in the WM stage have LTM associations, to assess potential modulations by object familiarity on the influence of experimental retro-cue manipulations during the WM stage.

Finally, our study focused specifically on visual LTM (Brady et al., 2013, 2008; Draschkow et al., 2019; Fukuda and Vogel, 2019; Schurgin, 2018; Standing, 1973) tested following a relatively short delay between our WM and LTM tasks (similar to the delays used in Fan and Turk-Browne, 2013; Hartshorne and Makovski, 2019; Jeanneret et al., 2023; LaRocque et al., 2015; Reaves et al., 2016; Sabo et al., 2024; Strunk et al., 2019). In future work, it will be important to extend our findings to other forms of LTM and to longer delays to see how generic and long-lasting the here reported effects are.

### General Conclusion

Our data make clear how attentional focusing during visual WM can instill a lasting effect on visual LTM that is driven by a benefit for the focused memorandum. We have shown how this benefit is supported by faster – not necessarily more – attentional deployment during WM. Our data open the door to future studies to extend our findings to other forms of memory, to test LTM after longer periods of time, and to causally manipulate the speed of attentional focusing during WM. Our findings may further prove of interest to future studies on patients with memory and/or attentional problems or in educational settings in which the transfer of information from WM to LTM is a central objective.

## Methods

### Participants

Twenty-five healthy human volunteers participated in Experiment 1, with ages ranging from 18 to 26 (16 female and 9 male; all right-handed). Sample size was set a-priori based on prior publications with comparable eye-tracking outcome variables (de Vries et al., 2023; Draschkow et al., 2022; Liu et al., 2022; van Ede et al., 2021; Van Ede et al., 2020; van Ede et al., 2019; Wang and van Ede, 2025, 2024). Experiment 2 extended Experiment 1 by adding an important additional condition with neutral cues, resulting in less trials per condition. To counteract having less trial per condition, we decided to double the sample size for Experiment 2. For Experiment 2, we recruited 50 participants, with ages ranging from 18 to 33 (34 female, 12 male, 4 non-binary; 46 right-handed, 1 left-handed, and 3 ambidextrous). Participant recruitment was performed independently for the two Experiments. Participants that had participated in Experiment 1 could not enrol in Experiment 2. The experimental procedures received approval from the Research Ethics Committee of the University. All participants provided written informed consent prior to participation, and received compensation of 11 dollars per hour, or course credits as reimbursement for their time.

### Sample Size Analysis

As explained in the Participants section, we set the sample size of the current experiments based on prior studies using similar outcomes measures. In addition, to determine the effect sizes that the current sample size would likely detect, we conducted a sensitivity power analysis for our sample sizes (N = 25 for Experiment 1 and N = 50 for Experiment 2) using the G*Power analysis software (Faul et al., 2009). Given the selected sample sizes and a target power of 0.8, we estimated the minimum detectable effect sizes to be *dz* = 0.512 for Experiment 1 and *dz* = 0.357 for Experiment 2.

### Task and Procedure

In both experiments, participants completed a two-stage WM-LTM task, with an attentional manipulation in the WM task. Our focus was on the influence of attentional prioritization during WM on subsequent LTM. In Experiment 1, all trials contained an informative retro-cue (presented during WM retention) that instructed which of two WM objects would be needed for the ensuing WM search task. In Experiment 2, we additionally included trials with a neutral retro-cue that enabled us to disentangle retro-cue benefits for cued (attended) WM objects from costs for uncued (unattended) WM objects. In both experiments, the WM task was followed by a surprise LTM test that enabled us to test the influence of prioritization during WM on subsequent visual LTM.

#### WM Task

The WM task is depicted in Figure 1a. Each trial started with an encoding display containing two images of real-world objects, each with a different color, presented for 500 ms. Memory objects were taken from the database in Brady et al 2013 (Brady et al., 2013) (for full details about the stimuli, see the “Stimuli” section below). The encoding display was followed by an initial delay period during which a central fixation dot remained on the screen for 1000 ms. Subsequently, the color of the central fixation dot changed to match one of the memorized objects for 500 ms, serving as the attentional retro-cue that indicated which memory object needed to be searched for after another delay of 1500 ms. Following this second delay, participants were presented with two greyscale objects on both sides of the screen. In trials with an informative retro-cue (all trials in Experiment 1), participants were always instructed to search exclusively for the previously cued memory object (i.e., the memorized object with the same color at encoding as the preceding cue) within the search array, and to indicate whether it was present or absent. In Experiment 2, we additionally included neutral trials. In these trials, the retro-cue color did not match either object in WM and participants were instead instructed to search for the presence of *either* memorized object during search.

The search array was displayed on the screen for 500 ms (matching the duration of the memory array). Participants used their fingers to press the “F” or “J” key on the keyboard, to indicate the presence or absence of the cued memory object, with the assignment of keys for “present” and “absent” fully counterbalanced across all participants. Following each response, feedback was provided on the fixation dot, with “1” denoting a correct response and “x” indicating an incorrect response. As mentioned above, the procedure for neutral-cue trials mirrored that of informative retro-cue trials, except that during the WM delay, the central fixation dot changed to a third color that did not match the color of either of the two memorized objects. In these trials, participants were instructed to indicate whether either memory object was present in the search array (or, alternatively, whether both were absent).

We carefully chose a search task for testing WM contents after the retro-cue. The search task with its critical target-absent trials enabled a vital control for equating sensory exposure between cued and uncued memory objects. Unlike in target-present trials, in target-absent trials, the cued/attended memory object never re-appeared at test. Accordingly, in these trials, any influence of retro-cueing on subsequent LTM cannot be attributed to differential sensory exposure between cued and uncued objects. Our main interest was thus in the target-absent trials where sensory exposure was equated between our conditions of interest. Considering this, the majority of trials (3 out of 4) were target-absent trials. The remaining trials with a target present in the search array were merely included to keep participants engaged in the WM task. Target-present and target-absent trials were randomly intermixed.

In both experiments, trials featuring different types of cues (retro-cue or neutral cue) and various testing conditions (present or absent) were intermixed within block for the WM task. Participants completed one WM block consisting of 192 and 216 trials for Experiment 1 and Experiment 2, respectively. In Experiment 1, all trials contained an informative retro-cue, while in Experiment 2, one third of the trials contained a neutral cue (72 trials) and the remaining 144 trials contained an informative cue. Because the ratio of the number of target-absent-at-search vs. target-present-at-search trials is 3:1, this resulted in 144 test-absent and 48 test-present trials in Experiment 1, and 162 test-absent and 54 test-present trials in Experiment 2.

#### LTM Task

Following completion of the WM task, participants were given a 10-minute break. During the 10-minute break, participants were free to do anything they wanted. Most participants chatted with the researcher and/or interacted with their mobile phones. Participants were not informed about the surprise LTM test until after the end of this 10-minute break. Therefore, there was no incentive to keep rehearsing the objects from the preceding WM task. This delay was set mainly for pragmatic reasons, and was in the same range as the delay used in complementary studies considering the relation between WM and LTM (Fan and Turk-Browne, 2013; Hartshorne and Makovski, 2019; Jeanneret et al., 2023; LaRocque et al., 2015; Reaves et al., 2016; Sabo et al., 2024; Strunk et al., 2019). When introducing the surprise LTM test, participants were instructed to report objects that they had seen at any point prior (during the WM task stage), regardless of whether they had been presented in the memory array or in the search array. They were informed that although they could not have known that these objects would be tested later, they might still retain some memories of them.

The LTM test is depicted in Figure 1b. During the LTM test, each test array consisted of two greyscale real-world objects positioned on the left and right sides of the screen, that required a 2 alternative-forced choice (2AFC) response. One of the two objects had always been presented during the WM stage (either as one of the encoded objects or as a filler in the search display). Another object during each 2AFC LTM test was always a completely novel object. Participants used their fingers to indicate whether the object presented on the left or right side of the screen was the one they had encountered in the preceding WM task (pressing “F” with the left finger if the old object was on the left, or “J” with the right finger if the old object was on the right, regardless of whether they were presented in the memory array or the search array). The locations of the old and new objects were fully counterbalanced across trials. After each response, feedback was provided on the fixation dot, with “1” indicating a correct response and “x” indicating an incorrect response.

In both experiments, participants completed one LTM test block consisting of 438 and 486 trials for Experiment 1 and Experiment 2, respectively. Within the LTM test, one out of seven trials (48 and 54 trials for E1 and E2, respectively) were images of fillers that only presented during WM search stage. The remaining trials were images of all objects presented during WM encoding stage (384 and 432 trials for E1 and E2, respectively). The order in which the objects would occur in the LTM test was independent of the order they would appear in the WM test.

#### Stimuli

The stimuli were presented using MATLAB (R2020a; MathWorks) and the Psychophysics Toolbox (version 3.0.16, Brainard, 1997) on a LED monitor. The monitor was a 23-inch (58.42-cm) screen operating at 240 Hz with a resolution of 1920 by 1080 pixels. Participants were seated 70 cm from the screen with their head resting on a chin rest to ensure stable eye-tracking.

Throughout the experiment, a white background (RGB = 255 255 255) was maintained, along with a grey fixation dot (0.8° visual angle, RGB = 192 192 192) presented in the center of the screen. The memory and search objects were real-world object pictures (obtained and modified from Brady et al., 2013) measuring 4° in size, presented at 6° eccentricity to central fixation. During WM encoding, the memory objects were displayed in two colors randomly chosen from a color pool comprising red (RGB = 158 73 76), green (RGB = 51 129 42), and purple (RGB = 82 88 172). The third color was used as the color of the neutral cue in Experiment 2. The two colors for the memory objects were consistent within each participant but randomized between participants to prevent any systematic sensory preference for a particular color. The memory objects were horizontally aligned to the fixation dot, with their colors and locations fully counterbalanced across trials. The objects in the search display were greyscale pictures (RGB = 128 128 128), presented at two out of four possible locations (top-left, top-right, bottom-right, bottom-right, 6° eccentricity to central fixation), with always one object on each side of the screen to balance visual inputs. The LTM test arrays comprised two greyscale objects (RGB = 128 128 128, size of 4°) presented on the left and right sides of the screen, horizontally aligned to the fixation dot (6° eccentricity to central fixation).

Color was exclusively used so that we could use a color change in the central fixation dot as a non-spatial cue to direct attention to either memory object during the WM task. The non-spatial nature of the cue was important to avoid sensory confounds when comparing our gaze data between cues directing attention to the left or right memory object. Color was thus used to cue the relevant object, but color itself was never tested; not during WM, nor during LTM.

All images came from the same database (Brady et al., 2013). In Experiment 1, the WM task featured 384 real-world object images used as memory objects (colored), while the search array included 336 separate filler objects (greyscale) randomly selected from the remaining images. The LTM recognition test contained 432 objects from the WM test (old objects, all in greyscale) and 432 novel objects. The 432 old objects comprised all 384 objects presented in the memory array (in greyscale during the LTM test), along with 48 fillers, with fillers randomly chosen from the total number of fillers used in the search array during the WM task. Novel objects were taken from a left-out subset of the images from the same object database. For convenience, we used the same subset of images as memory objects, filler objects, and novel objects for all participants. Within the pool of memory objects, the designation into the different attentional conditions (cued/uncued/neutral) was randomized, and was thus different for each participant. Experiment 2 utilized 432 colored images for the memory array, 378 fillers for the search array during WM task, and 486 old and 486 novel objects for the LTM recognition test. The 486 old objects included all 432 objects presented in the memory array in greyscale, along with 54 (1/7 of all fillers) randomly chosen objects from the fillers used in the search array during the WM task.

### Analysis of behavioral data

We analysed both response accuracy and response time (RT), for both the WM and the LTM task. Our focus was on performance in the LTM 2AFC recognition task, as a function of objects’ attentional status during WM (cued and uncued in Experiment 1; cued, uncued and neutral in Experiment 2). We separately considered objects from target-present and target-absent WM-search trials, with our focus on target-absent-during-search trials in which sensory exposure was equated between attention conditions (as detailed above). For completeness, we also included ‘search-only’ objects (i.e., objects that served as fillers in the search arrays) when considering LTM recognition of objects encountered in the WM task.

### Eye-tracking acquisition and pre-processing

The eye tracker (EyeLink 1000, SR Research SR) was positioned approximately 5 cm in front of the monitor on a table, approximately 65 cm away from the participants’ eyes. Horizontal and vertical gaze positions were continuously recorded for a single eye at a sampling rate of 1000 Hz. Prior to recording, the eye tracker underwent calibration using the built-in calibration and validation protocols provided by the Eye-Link software. Participants were instructed to keep their chin stable on the chin rest for the entire block following calibration.

For the offline analyses, eye-tracking data were converted from the .edf to the .asc format and imported into MATLAB using Fieldtrip (Oostenveld et al., 2011). Blinks were identified by detecting clusters of zeros in the eye-tracking data. Subsequently, data from 100 ms before to 100 ms after the detected blink clusters were marked as Not-a-Number (NaN) to be excluded from further analysis. Following blink rejection, data were epoched relative to the onset of the cue in the WM task.

### Saccade detection

We utilized a velocity-based approach for saccade detection (as in: de Vries et al., 2023; Liu et al., 2023; Liu et al., 2022) that was developed building on closely-related velocity-based methods for saccade detection (e.g., Engbert and Kliegl, 2003). Because the memory objects in our experiment were always presented horizontally at center-left and center-right positions, our saccade detection targeted the horizontal channel of the eye data. This approach has previously demonstrated to be effective for capturing directional biases in saccades during internal attention shifts, as expanded on in Liu et al., 2022.

We computed gaze velocity by measuring the distance between gaze position in consecutive samples. To improve accuracy and reduce noise, we applied temporal smoothing to the velocity using a Gaussian-weighted moving average filter with a 7-ms sliding window, utilizing the “smoothdata” function in MATLAB. Subsequently, we identified the onset of a saccade as the first sample where the velocity exceeded a trial-based threshold set at 5 times the median velocity. To prevent multiple classifications of the same saccade, we imposed a minimum delay of 100 ms between successive saccades. Saccade magnitude and direction were determined by evaluating the difference between pre-saccade gaze position (from -50 to 0 ms before threshold crossing) and post-saccade gaze position (from 50 to 100 ms after threshold crossing). Each identified saccade was then classified as “toward” or “away” based on its direction (left/right) and the side of the cued memory object (left/right). “Toward” denoted a saccade in the direction of the memorized location of the cued object, while “away” indicated a saccade in the opposite direction.

Following saccade identification and categorization, we analyzed the time courses of saccade rates (in Hz) using a sliding time window of 50 ms, progressing in 1 ms increments. For our primary analyses, we focused directly on the spatial bias in saccades, calculated as the differences (in Hz) between toward and away saccades.

Finally, to further characterize the saccade bias, we computed the saccadic bias as a function of saccade size (as in: de Vries et al., 2023; Liu et al., 2023; Liu et al., 2022). For this, we quantified the saccade bias (toward vs. away) iteratively for saccades falling within specific saccade-size bins. We used size bins of 0.5 degrees, that were advanced in steps of 0.1 degrees.

### Subsequent memory effect

Our primary interest was to link attentional deployment during WM to subsequent changes in LTM performance. To this end, capitalized on the classic subsequent-memory effect (SME: Bartsch et al., 2018; Khader et al., 2007, 2010; LaRocque et al., 2015; Ranganath et al., 2005; Voss and Paller, 2009), and applied it to our gaze signature of attentional allocation during WM. We focused on cued memory objects and split the WM trials depending on whether these cued objects were later remembered or later forgotten during the LTM test.

Having established a subsequent memory effect in our data, we wished to rule out obvious “third factors” that could have supported our effect. First, we considered object memorability itself. Some objects may be more memorable, and these objects may render different attentional allocation during WM. To rule this out, we also sorted our WM gaze data based on object memorability, using a median split. We did this twice, following two different ways to quantify object memorability. First, we estimated the memorability of all objects by sorting them by the overall correct identification rate during the LTM test, collapsed across our full sample of participants and combining both experiments for maximum sensitivity. In addition to our first approach, where we defined object memorability empirically based on our own LTM performance data, we also quantified object memorability using a state-of-the-art machine learning model for predicting the intrinsic memorability of an image – ResMem (Needell and Bainbridge, 2022). Next to object memorability, we also considered time-on-task as another factor that may underlie our observed subsequent memory effect. For this, we simply split our WM data into ‘early’ and ‘late’ trials (i.e., first half vs. second half of WM trials), to compare our gaze data of interest as a function of time on task.

The use of median splits was motivated by the fact that saccades that make up our gaze bias marker are themselves discrete short-lived events that yield a continuous gaze-bias time course measure only after averaging across multiple trials. Moreover, our gaze bias is calculated as the rate of toward vs. away saccades, which further necessitates aggregating multiple trials (trials in which saccades were made toward vs. away from the cued memory object). In addition, the median-split approach further enabled us to visualize the relevant gaze-bias time courses separately for each split, which is critical for disambiguating relevant variability in amplitude from relevant variability in latency. Please also note that, in addition to this binary categorization of the relevant data, for completeness, we also performed quartile splits.

### Saccade bias latency analysis

To evaluate condition differences in the timing of the spatial saccade bias following the cue, we utilized a simplified jackknife method (as described in Smulders, 2010). Specifically, we compared the onset latencies of the spatial saccade bias after the cue in the WM task between later-remembered and later-forgotten objects, as derived from responses on the LTM task. The onset latency was determined by the time at which the amplitude reached 50% of the peak value.

### Statistical analysis

For the performance data, our main focus was on accuracy and RT in the LTM task, as a function of attentional status of the objects during the preceding WM task. We conducted pre-planned paired-sample t-tests between conditions with different cueing and probing types during the WM task on the LTM task. Our focus was on WM objects in trials in which the cued object was absent in the search array (in which sensory exposure was equated between cued, uncued, and neutral objects). For completeness, we also considered objects in target-present-during-search trials, and objects that only appeared at test as fillers in the search display. To account for multiple comparisons, all reported p-values were Bonferroni corrected. In addition, in Experiment 2 we also compared performance on the WM task between trials with informative vs. neutral cues by using a two-way repeated measures ANOVA with within-subject factors of Cue Type (informative cue vs. neutral cue) and Search Type (present vs. absent) on WM accuracy and RTs (in Experiment 1 all trials contained an informative retro-cue, so this comparison could not be made).

To analyze the temporal patterns of spatial modulations in eye movements (saccade bias), we utilized a cluster-based permutation approach (Maris and Oostenveld, 2007) implemented with the ‘ft_timelockstatistics’ function from the Fieldtrip toolbox, employing the ‘montecarlo’ method. This methodology is particularly well-suited for assessing the consistency of data patterns across multiple adjacent data points (effectively circumventing the issue of multiple comparisons by evaluating the full time-course under a single permutation distribution of the largest cluster). We conducted 10,000 permutations to generate the permutation distribution of the largest cluster that could occur by chance. Clusters were identified using Fieldtrip’s default settings, which involve grouping temporally adjacent data points with the same sign that were statistically significant in a mass univariate t-test at a two-sided alpha level of 0.05. The size of each cluster was defined as the sum of all t values within that cluster.

For the saccade bias latency comparison between later-remembered vs. later-forgotten objects, we used standard paired-samples t-tests to compare the latency estimates that we obtained from the employed jackknife methods (Smulders, 2010). Where relevant, we supplemented our findings with Bayesian Factor analyses to quantify the amount of evidence in favor of the null hypothesis of no difference between conditions. This was conducted using open-source software JASP (version 0.18.3) with the default settings.

## Acknowledgments

This research was supported by an ERC Starting Grant from the European Research Council (MEMTICIPATION, 850636) and an NWO Vidi Grant by the Dutch Research Council (grant number 14721) to F.v.E.

## Supplemental Materials

**Figure S1.**
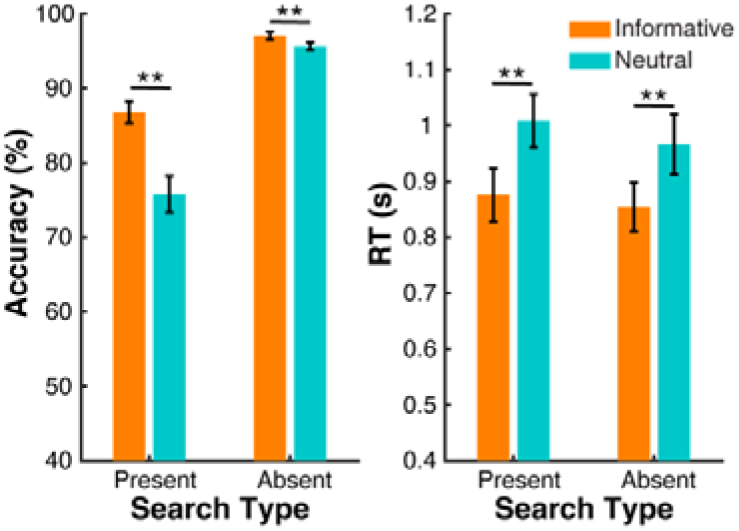
Attentional cueing effects on WM performance in Experiment 2. The WM search performance for objects with different cue types in Experiment 2, where we included both informative and neutral cues. In both target-present-during-search and target-absent-during-search trials, the search performance (accuracy-left panel and RT-right panel) for the informatively cued objects outperformed trials with neutral cues. Orange and blue bars represent informative and neutral cue trials, respectively. Error bars represent ±1SEM. *, ** represent significance level p < 0.05 and p < 0.01 after Bonferroni correction.

**Table S1.** Statistics for LTM recognition performance. *, *** represent significance level *p* < 0.05 and *p* < 0.001 after Bonferroni correction. Conditions with bold font represent our key comparisons.

| Experiment 1 |  |  |  | Experiment 2 |  |  |  |
| --- | --- | --- | --- | --- | --- | --- | --- |
| LTM recognition accuracy |  |  |  | LTM recognition accuracy |  |  |  |
| | | $t_{(24)}$ | $p$ | | | $t_{(49)}$ | $p$ |
| Cued-present | Cued-absent | 2.747* | .046 | Cued-present | Cued-absent | 2.671 | .121 |
|  | Uncued-absent | 7.461*** | <.001 |  | Uncued-absent | 6.939*** | <.001 |
|  | Searched-only | 20.039*** | <.001 |  | Neutral-present | 1.277 | 1.000 |
|  |  |  |  |  | Neutral-absent | 6.722*** | <.001 |
|  |  |  |  |  | Searched-only | 21.742*** | <.001 |
| **Cued-absent** | **Uncued-absent** | **4.714\*\*\*** | **<.001** | **Cued-absent** | **Uncued-absent** | **4.268\*\*\*** | **<.001** |
|  | Searched-only | 17.292*** | <.001 |  | Neutral-present | -1.394 | 1.000 |
|  |  |  |  |  | **Neutral-absent** | **4.052\*\*\*** | **<.001** |
|  |  |  |  |  | Searched-only | 19.071*** | <.001 |
| Uncued-absent | Searched-only | 122.578*** | <.001 | **Uncued-absent** | Neutral-present | -5.661*** | <.001 |
|  |  |  |  |  | **Neutral-absent** | **-.216** | **1.000** |
|  |  |  |  |  | Searched-only | 14.803*** | <.001 |
|  |  |  |  | Neutral-present | Neutral-absent | 5.445*** | <.001 |
|  |  |  |  |  | Searched-only | 20.464*** | <.001 |
|  |  |  |  | Neutral-absent | Searched-only | 15.019*** | <.001 |

**Figure S2.**
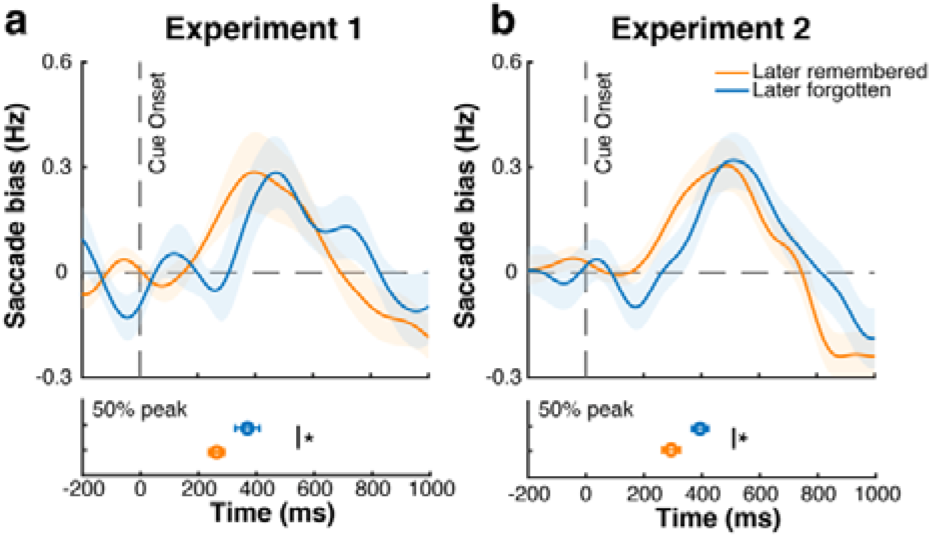
Faster attentional deployment in WM for later-remembered objects cannot be attributed to uneven trial numbers between these trial-classes. **a**) Top panel: spatial saccade bias (toward minus away) after the retro-cue during the WM task, for later-remembered and later-forgotten objects in Experiment 1, with trials in the later-remembered condition sub-sampled to the same trial number as in the later-forgotten condition. Bottom panel: latency of the saccade bias, defined as the 50% of the peak value. The error bars represent standard errors obtained using a jackknife approach. **b**) Same as in panel a, but for Experiment 2. Shading and error bars represent ±1SEM. * represent significance level *p* < 0.05 after Bonferroni correction.

**Figure S3.**
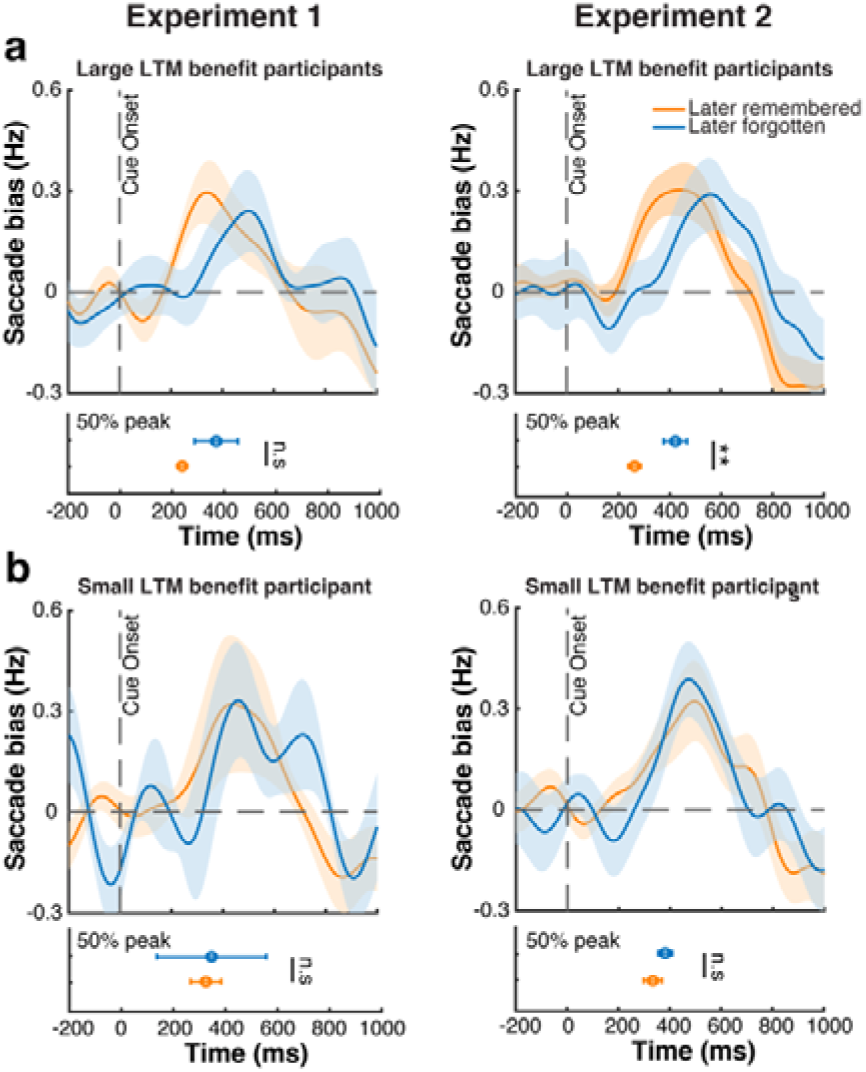
The latency of attentional deployment in WM predicts subsequent LTM particularly for individuals with larger LTM benefit across both experiments. **a**) Gaze bias for later-remembered and later-forgotten trials in the subset of participants with relatively little benefit of cueing during WM on subsequent LTM across both experiments (left panel – Experiment 1, right panel – Experiment 2). **b**) Gaze bias for later-remembered and later-forgotten trials in the subset of participants with relatively large benefit of cueing during WM on subsequent LTM across both experiments (left panel – Experiment 1, right panel – Experiment 2). Shading and error bars represent ±1SEM. *, ** represent significance level *p* < 0.05 and *p* < 0.01 after Bonferroni correction.

**Figure S4.**
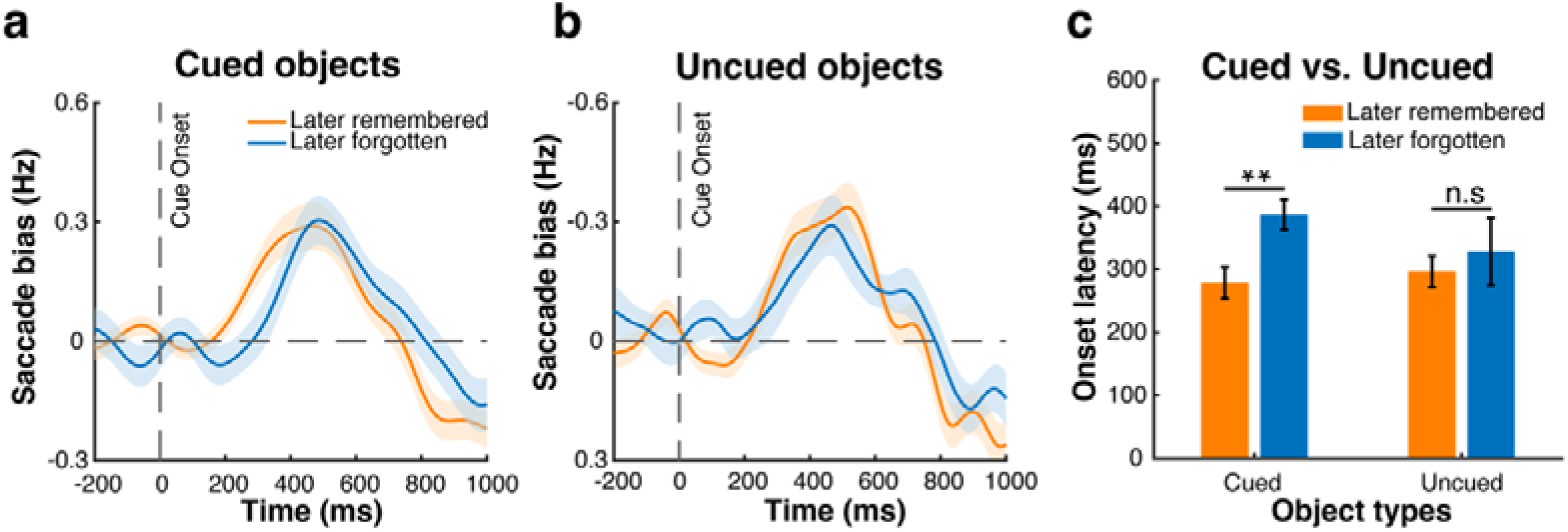
Saccade bias as a function of subsequent memory performance for cued and uncued objects. **a**) Spatial saccade bias (toward minus away from the cued object) after the retrocue during the WM task, separately for later-remembered and later-forgotten cued objects (Experiment 1 and Experiment 2 combined). **b**) Spatial saccade bias (toward minus away from the uncued object) after the retro-cue during the WM task, separately for later-remembered and later-forgotten uncued objects (Experiment 1 and Experiment 2 combined). **c**) Bar-graph of gaze-bias onset latency for subsequent remembered and subsequent forgotten cued and uncued objects. Shading and error bars represent ±1SEM. ** represents significance level *p* < 0.01 (*t*_(74)_ = -3.349, *p* = 0.001) after Bonferroni correction, n.s represents not significant (*t*_(74)_ = -0.562, *p* = 0.576).

**Figure S5.**
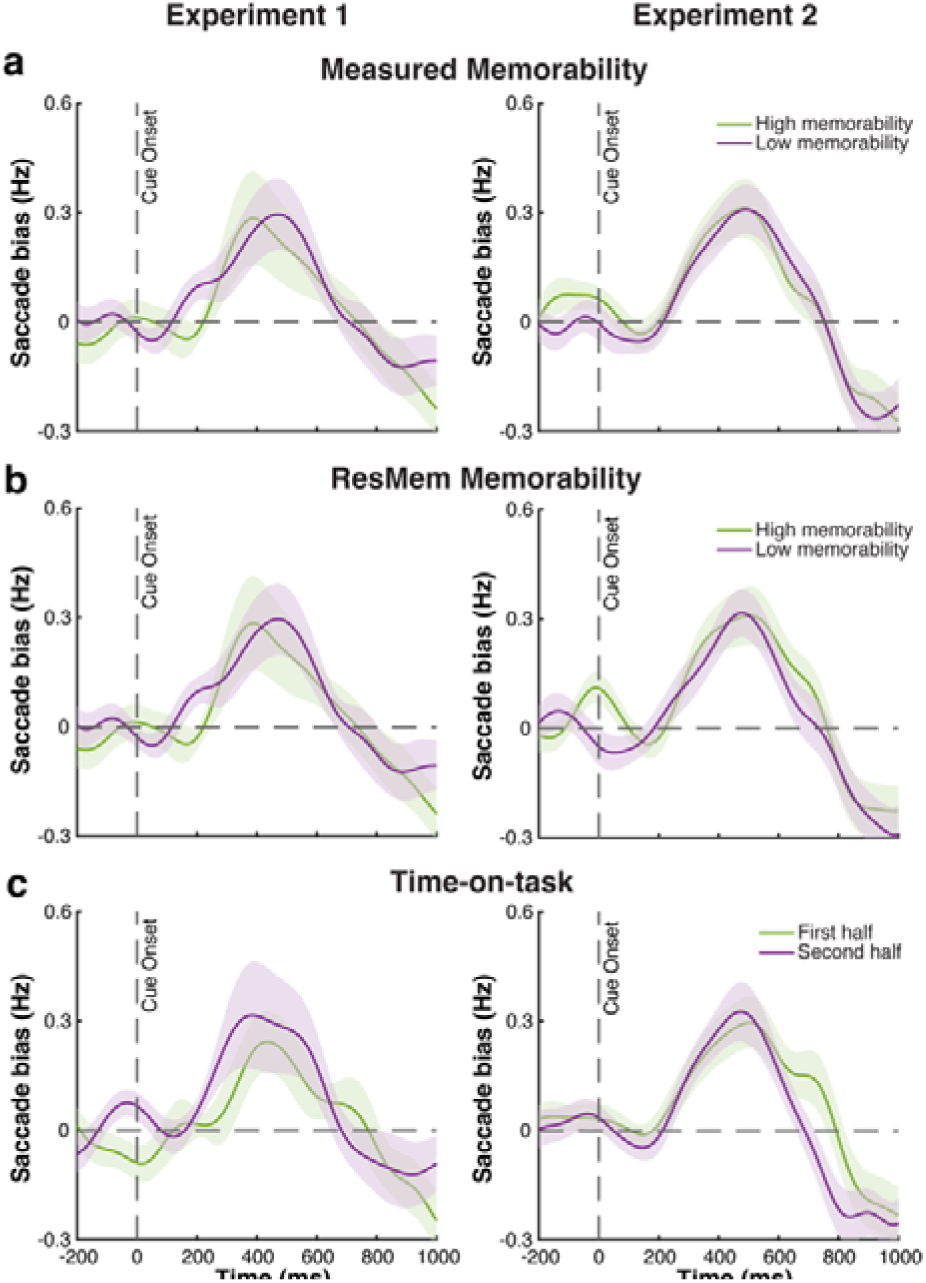
The observed relation between the timing of attentional deployment and subsequent LTM is not driven by object memorability nor by time-on-task across both experiments. **a**) Spatial saccade bias after the informative cue during the WM task for cued objects with high (green) and low (purple) general memorability across both experiments (left panel – Experiment 1, right panel – Experiment 2). Memorability was estimated from LTM performance from the current experiments, and objects were sorted using a median split. **b**) As panel a, but quantifying object memorability using ResMem (Needell and Bainbridge, 2022). **c**) Spatial saccade bias for objects presented in the first-half (green lines) vs. objects presented in the second-half (purple lines) of the WM session across both experiments (left panel – Experiment 1, right panel – Experiment 2). Bottom panels and all other conventions as in preceding figures. Shading areas represent ±1SEM.

**Figure S6.**
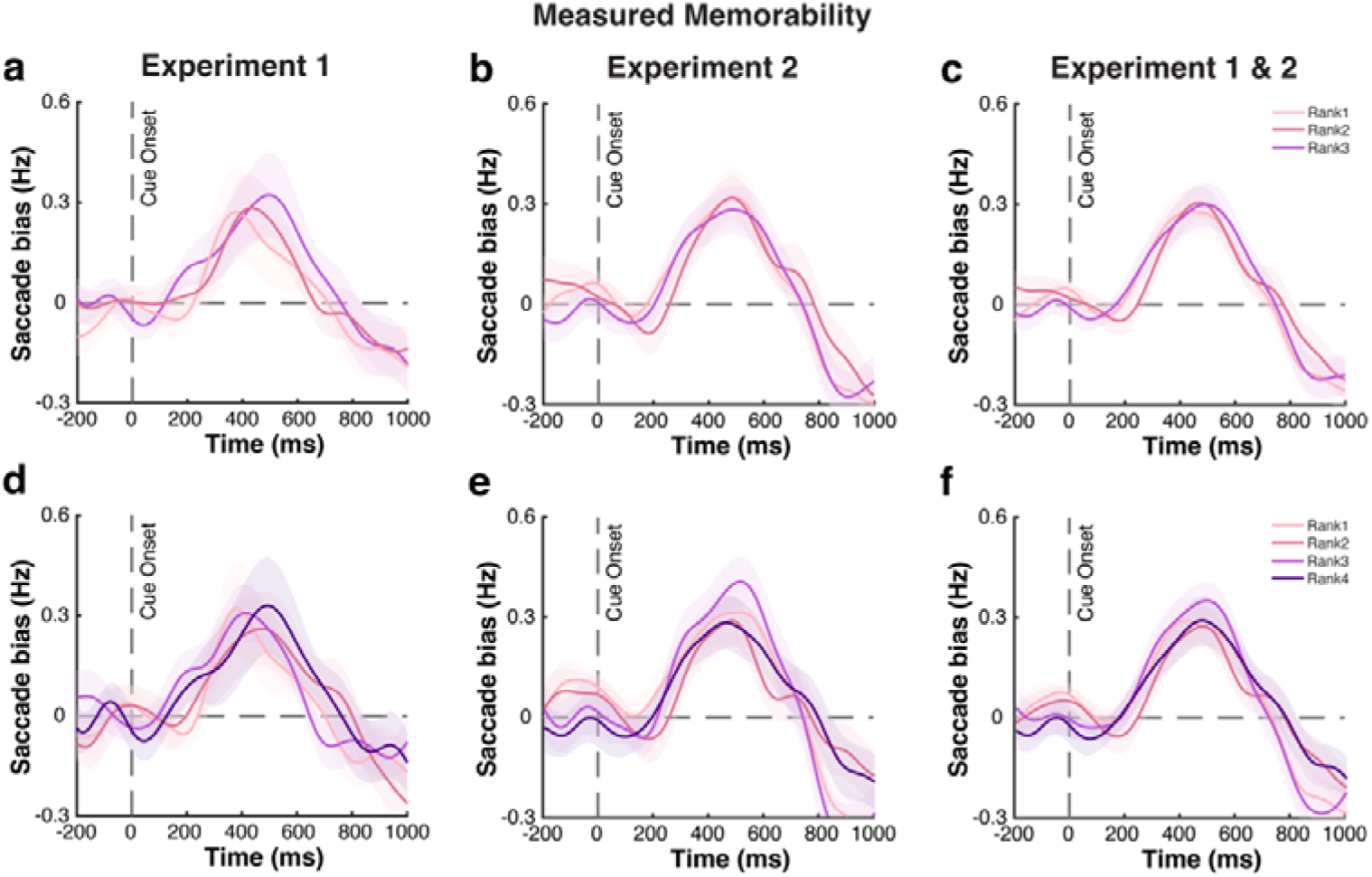
The observed relation between the timing of attentional deployment and subsequent LTM is not driven by object memorability (tertile- and quartile-split). Top panels: **a**) Spatial saccade bias after the informative cue during the WM task for cued objects with different ranks of general memorability (tertile-split: rank 1 to rank 3, from high to low memorability) in Experiment 1. **b**) & **c**) same results for Experiment 2, and the merged dataset of both experiments. **Bottom panels: d**) Spatial saccade bias after the informative cue during the WM task for cued objects with different ranks of general memorability (quartile-split: rank 1 to rank 4, from high to low memorability) in Experiment 1. **e**) & **f**) same results for Experiment 2, and the merged dataset of both experiments. Memorability was estimated from LTM performance from the current experiments, and objects were sorted using a quartile-split method. Shading areas represent ±1SEM.

For the control analysis of Memorability, instead of exclusively relying on a median split, we additionally split all objects into three and four groups with different ranks of memorability from high to low, respectively. One-way repeated measures ANOVAs with within-subject factor of Memorability (tertile-split: rank 1 vs. rank 2 vs. rank 3; quartile-split: rank 1 vs. rank 2 vs. rank 3 vs. rank 4) on the onset latency of saccade bias showed no statistically significant difference between them across Experiment 1 (tertile-split: *F*(2,48) = 0.299, *p* = 0.743, η*²p* = 0.012; quartile-split: *F*(3,72) = 0.218, *p* = 0.884, η*²p* = 0.009), Experiment 2 (tertile-split: *F*(2,98) = 0.389, *p* = 0.679, η*²p* = 0.008; quartile-split: *F*(3,147) = 0.698, *p* = 0.555, η*²p* = 0.041), nor for the merged dataset containing the data from both experiments (tertile-split: *F*(2,148) = 0.804, *p* = 0.450, η*²p* = 0.011; quartile-split: *F*(3,222) = 0.652, *p* = 0.583, η*²p* = 0.009). These results corroborate that the memorability of the pictures does not have a significant influence on the speed of saccade bias after the cue, no matter whether two (median), three (tertile), or four (quartile) bins were used for splitting the trials.

**Figure S7.**
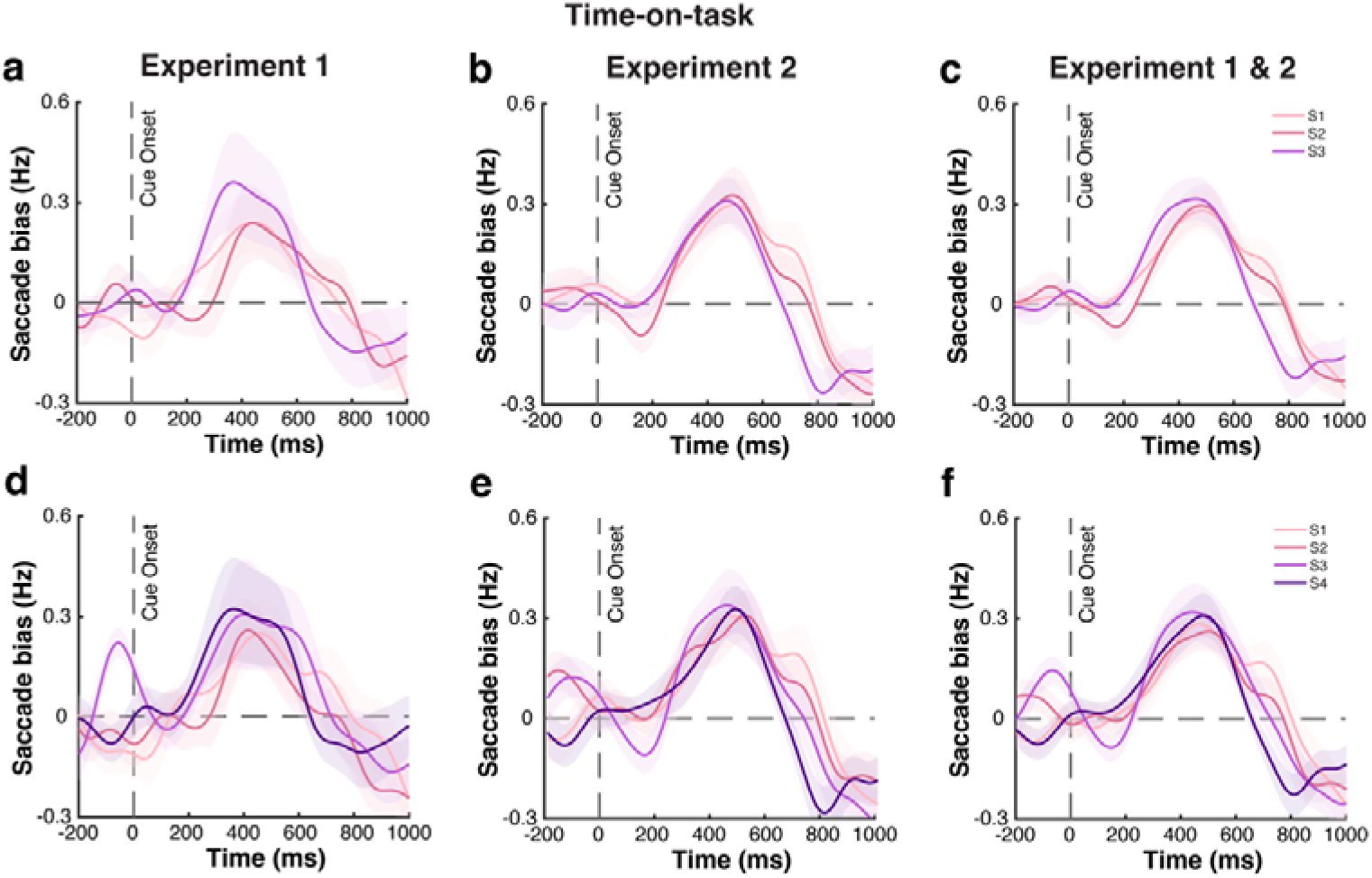
The observed relation between the timing of attentional deployment and subsequent LTM is not driven by time-on-task (tertile-and quartile-split). Top panels: **a**) Spatial saccade bias after the informative cue during the WM task for cued objects with different time-on-task (tertile-split: session 1 to session 3, from early to late) in Experiment 1. **b**) & **c**) same results for Experiment 2, and the merged dataset of both experiments. **Bottom panels: d**) Spatial saccade bias after the informative cue during the WM task for cued objects with different time-on-task (quartile-split: session 1 to session 4, from early to late) in Experiment 1. **e**) & **f**) same results for Experiment 2, and the merged dataset of both experiments. Time-on-task were sorted using a quartile-split method. Shading areas represent ±1SEM.

For the control analysis of Time-on-task, we also split all trials into three and four sessions, respectively (from early to late). One-way repeated measures ANOVAs with within-subject factor of Time-on-task (tertile-split: session 1 vs. session 2 vs. session 3; quartile-split: session 1 vs. session 2 vs. session 3 vs. session 4) on the onset latency of saccade bias showed no statistically significant difference between them across Experiment 1 (tertile-split: *F*(2,48) = 2.050, *p* = 0.140, η*²p* = 0.079; quartile-split: *F*(3,72) = 0.113, *p* = 0.952, η*²p* = 0.005), Experiment 2 (tertile-split: *F*(2,98) = 0.156, *p* = 0.856, η*²p* = 0.003; quartile-split: *F*(3,147) = 1.517, *p* = 0.213, η*²p* = 0.030), nor for the merged dataset containing the data from both experiments (tertile-split: *F*(2,148) = 0.986, *p* = 0.376, η*²p* = 0.013; quartile-split: *F*(3,222) = 0.867, *p* = 0.459, η*²p* = 0.012). These results again corroborate that time-on-task itself does not have a significant influence on the speed of saccade bias after the cue, no matter whether two (median), three (tertile), or four (quartile) bins were used for splitting the trials.

## Notes

### Competing Interest Statement

The authors have declared no competing interest.

### Summary of Updates

1. Control analyses addressing the uneven number of trials between later-remembered and later-forgotten objects. 2. Control analyses examining potential confounding factors using more sophisticated trial-grouping methods, for example, added quarter-split analyses in addition to the median-split. 3. Updated statistical analyses, for example, Bayes factor analyses, and more complete statistical power reporting are included.

